# CaMKII autophosphorylation but not downstream kinase activity is required for synaptic memory

**DOI:** 10.1101/2023.08.25.554912

**Authors:** Xiumin Chen, Qixu Cai, Jing Zhou, Samuel J. Pleasure, Howard Schulman, Mingjie Zhang, Roger A. Nicoll

## Abstract

CaMKII plays a critical role in long-term potentiation (LTP), a well-established model for learning and memory through the enhancement of synaptic transmission. Biochemical studies indicate that CaMKII catalyzes a phosphotransferase (kinase) reaction of both itself (autophosphorylation) and of multiple downstream target proteins. However, whether either type of phosphorylation plays any role in the synaptic enhancing action of CaMKII remains hotly contested. We have designed a series of experiments to define the minimal requirements for the synaptic enhancement by CaMKII. We find that autophosphorylation of T286 and further binding of CaMKII to the GluN2B subunit are required both for initiating LTP and for its maintenance (synaptic memory). Once bound to the NMDA receptor, the synaptic action of CaMKII occurs in the absence of kinase activity. Thus, autophosphorylation, together with binding to the GluN2B subunit, are the only two requirements for CaMKII in synaptic memory.

## Introduction

The alpha isoform of Ca^2+^/calmodulin (CaM) dependent kinase II enzyme (referred to throughout as CaMKII) has emerged as a central player in synaptic plasticity (Bayer and Schulman, 2019; Bhattacharyya et al., 2020; Hell, 2014; Lisman and Raghavachari, 2015; Nicoll and Schulman, 2023; Shonesy et al., 2014; Yasuda et al., 2022). While activation of CaMKII is required for the induction of NMDA-dependent long-term potentiation (LTP), an established cellular model for learning and memory, the precise steps following its activation remains unclear. CaMKII is activated by an NMDA receptor-dependent rise in Ca^2+^ which together with CaM displaces an inhibitory segment of the kinase, converting the inactive closed conformation into an active open conformation. In the open conformation, CaMKII autophosphorylates T286 in the inhibitory segment, disables the inhibitory segment and uncovers the binding sites for its substrates. The open conformation of CaMKII can be generated experimentally by use of the phosphomimic mutation T286D. CaMKII binds (and phosphorylates) target substrate proteins as well as binding anchoring proteins such as the GluN2B subunit of the NMDAR.

The relative importance of T286 autophosphorylation, phosphorylation of downstream target proteins and the binding to GluN2B remain hotly debated (Bayer and Schulman, 2019; Nicoll and Schulman, 2023; Yasuda *et al*., 2022). Depending on the literature one can select between two extreme scenarios can be proposed: enzymatic and structural. In the enzymatic scenario LTP is maintained by T286 autophosphorylation and the phosphorylation of target substrate proteins. In the structural scenario LTP is maintained by the binding of CaMKII to GluN2B independent of any enzymatic activity toward target proteins.

To disambiguate among these scenarios, we determined the minimal requirements for the ability of CaMKII to enhance synaptic transmission and its role in the maintenance of LTP. We establish that T286 autophosphorylation is, indeed, required both for the induction and maintenance of LTP. Remarkably, a CaMKII kinase dead mutant (D135N), maintained in an open conformation by the additional T286D mutation, is fully capable of enhancing synaptic transmission. CaMKII synaptic function requires the binding to the GluN2B subunit of the NMDAR. We conclude that the only required kinase role of CaMKII in LTP is its autophosphorylation of T286, which, by enabling and stabilizing the CaMKII/GluN2B complex, maintains synaptic memory independent of phosphorylation of downstream targets.

## Results

### CaMKII mediated synaptic enhancement is independent of substrate protein phosphorylation

We first expressed a phosphomimic mutant of CaMKII (Incontro et al., 2018; Pi et al., 2010) which maintains the enzyme in an open conformation, referred to as constitutively active CaMKII (CA CaMKII). Simultaneous whole cell recordings were made from a control cell and a transfected cell and the responses from a common synaptic input compared (**Fig. 1A**). When overexpressed for a short period of time (1-5 days), referred to as short overexpression (short OE), CA CaMKII causes an approximately 3-fold enhancement in AMPAR responses (**Fig. 1B and D**, n = 15), in general agreement with previous studies (Hayashi et al., 2000; Incontro *et al*., 2018; Lledo et al., 1995; Pettit et al., 1994; Pi *et al*., 2010; Poncer et al., 2002; Shirke and Malinow, 1997; Wu et al., 1996). Importantly, CA CaMKII has no effect on the NMDAR responses (**Fig. 1C and D**, n = 15), thus mimicking LTP.

**Figure 1.**
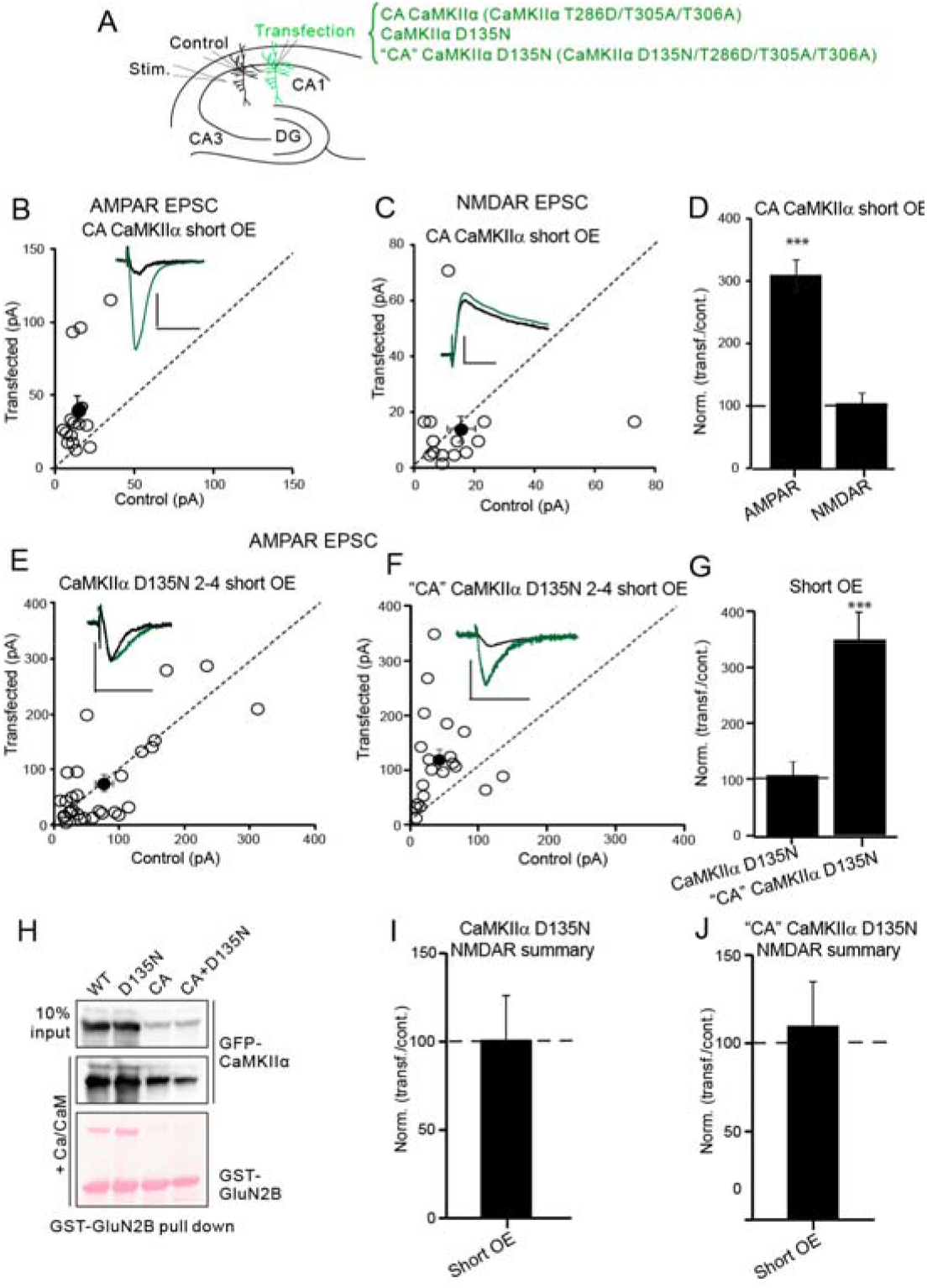
CaMKII mediated synaptic enhancement is independent of substrate protein phosphorylation. **A** Schematic diagram showing the electrophysiological approaches and three transfected CaMKII mutate constructs (Control represents the wild type, un-transfected neurons). **B** Scatterplots showing amplitudes of AMPAR EPSCs for single pairs (open circles) of control cell and cells overexpressing CA CaMKII 1-5 days (short OE) (n = 15 pairs). Filled circle indicate mean ± SEM. (Control = 15 ± 2; CA CaMKII short OE = 41 ± 8.8 p < 0.0001). **C** Scatterplots showing amplitudes of NMDAR EPSCs for single pairs (open circles) of control cells and cells transfected with CA CaMKII 1-5 days (short OE) (n = 15 pairs). Filled circles indicate mean ± SEM. (Control= 15 ± 5; CA CaMKII short OE = 14 ± 5, p = 0.9). **D** Bar graph of ratios normalized to control (%) summarizing the mean ± SEM of AMPAR and NMDAR EPSCs of values represented in B (314 ± 26, p < 0.0001) and C (118 ± 20, p =0.98). **E, F** Scatterplots showing amplitudes of AMPAR EPSCs for single pairs (open circles) of control and overexpressing cells of CaMKII D135N 2-4 days (short OE) (**E**, n = 30 pairs), and “CA” CaMKII D135N 2-4 days (short OE) (**F**, n = 25 pairs). Filled circle indicate mean ± SEM. (**E**, Control = 76.1 ± 13.9; CaMKII D135N short OE = 75.9 ± 15 p =0.98; **F**, Control = 42.5 ± 8; “CA” CaMKII D135N short OE = 120 ± 18.2 p < 0.0001). **G** Bar graph of ratios normalized to control (%) summarizing the mean ± SEM of AMPAR EPSCs of values represented in E (120 ± 23, p =0.6) and **F** (350 ± 50, p < 0.0001). **H** GST-GluN2B pull-down. In presence of Ca^2+^/CaM, CaMKIIα wt, CaMKIIα D135N, CA CaMKIIα and CA CaMKIIα D135N can bind to GluN2B. **I** Bar graph of ratios normalized to control (%) summarizing the mean ± SEM of NMDAR EPSCs of CaMKII D135N short OE (101 ± 25, p =0.6). **J** Bar graph of ratios normalized to control (%) summarizing the mean ± SEM of NMDAR EPSCs of “CA” CaMKII D135N long OE (95 ± 15, p =0.7). Raw amplitude data from dual cell recordings were analyzed using Wilcoxon signed rank test (p values indicated above). Normalized data were analyzed using a one-way ANOVA followed by the Mann−Whitney test. Scale bars: 30 ms, 50 pA.

Does the enhancement require enzymatic activity? To answer this question, we used CaMKII (D135N), a mutation that blocks the catalytic activity by eliminating the catalytic base D135 without perturbing kinase structure (Ozden et al., 2022) (**Fig. S1F**). Importantly this mutation does not impact the binding to GluN2B (**Fig. 1H**). Short OE of CaMKII (D135N) on its own had no effect on AMPAR responses (**Fig. 1E**, n =30). Remarkably, however, when this mutation was inserted into CA CaMKII (referred to as “CA” CaMKII (D135N) with the quotation marks to indicate that while in the constitutively open conformation it is not catalytically active), this kinase dead construct enhanced synaptic responses to the same degree as CA CaMKII alone either for short OE (**Fig. 1F**, n =25) or for 10-14 days (long OE) (**Fig. S1A-C**, n = 27). These findings are provocative because they indicate that phosphorylation of the numerous potential downstream targets is not necessary for the enhancing action of CaMKII.

The results with the kinase dead “CA” CaMKII (D135N) mutation suggest that the enhancing action of CaMKII does not require any downstream phosphorylation event. However, it is conceivable that the expressed “CA” CaMKII (D135N) could exchange either by subunit exchange (Stratton et al., 2014) or interholoenzyme phosphorylation (Lucic et al., 2023) with the constitutively active endogenous CaMKIIα present at synapses (Incontro *et al*., 2018; Lee et al., 2022; Sanhueza et al., 2011; Tao et al., 2021). To test this possibility, we made slices from the CaMKIIα KO mouse. Overxpression of wt CaMKII had no effect (**Fig. 2A** and **G**, n = 20). On the other hand, overexpression of CA CaMKII (**Fig. 2B** and **G**, n = 25) and “CA” CaMKII (D135N) (**Fig. 2C** and **G**, n = 22), still enhances synaptic transmission by 3-fold, confirming the conclusion that phosphorylation is not required for the enhancing action of expressed CA CaMKII. No change in the NMDAR response was observed with any of these manipulations (**Fig. 2D-F** and **H**).

**Figure 2.**
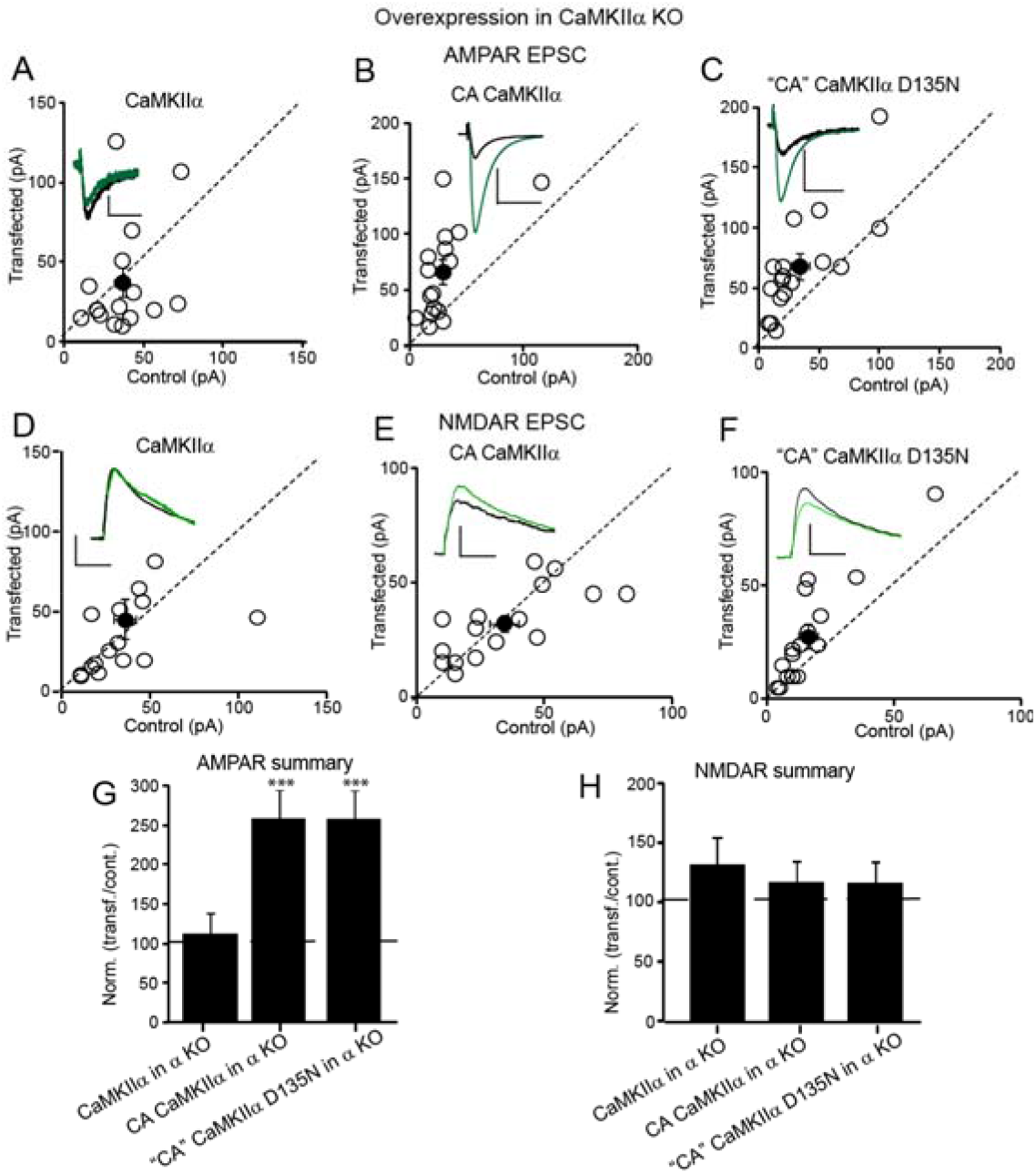
CaMKII mediated synaptic enhancement is independent of substrate protein phosphorylation even in the absence of endogenous CaMKIIα. **A-C** Scatterplots showing amplitudes of AMPAR EPSCs for single pairs (open circles) of control cells and cells expressing CaMKII wt for 2-3 days (short OE) (A, n = 20 pairs), CA CaMKII for 2-3 days (short OE) (**B**, n = 25 pairs), and “CA” CaMKII D135N for 2-3 days (short OE) (**C**, n = 22 pairs) all in CaMKIIα KO mice slices. Filled circle indicate mean ± SEM. (**A**, Control = 36.3 ± 4.7; CaMKII wt = 37.1 ± 9 p = 0.9; **B**, Control = 29.4 ± 6.1; CA CaMKII = 66.1 ± 10.6 p < 0.001; **C**, Control = 34.1 ± 7.5; “CA” CaMKII D135N = 68.2 ± 10.7 p < 0.001). **D-F** Scatterplots showing amplitudes of NMDAR EPSCs for single pairs (open circles) of control cells and cells expressing CaMKII wt for 2-3 days (short OE) (**D**, n = 20 pairs), CA CaMKII for 2-3 days (short OE) (**E**, n = 25 pairs), and “CA” CaMKII D135N for 2-3 days (short OE) (**F**, n = 22 pairs) all in CaMKIIα KO mice slices. Filled circle indicate mean ± SEM. (**D**, Control = 35.4 ± 6.4; CaMKII wt = 45.2 ± 12.5 p =0.6; **E**, Control = 34.3 ± 5.5; CA CaMKII = 32.1 ± 3.8 p =0.9; **F**, Control = 16.3 ± 3.7; “CA” CaMKII D135N = 27.6 ± 5.7 p =0.3). **G** Bar graph of ratios normalized to control (%) summarizing the mean ± SEM of AMPAR EPSCs of values represented in **A** (113 ± 25, p =0.8); **B** (263 ± 37, p < 0.001) and **C** (257 ± 33, p < 0.001). **H** Bar graph of ratios normalized to control (%) summarizing the mean ± SEM of NMDAR EPSCs of values represented in **D** (130 ± 23, p =0.1); **E** (117 ± 18, p =0.9) and **F** (121 ± 19, p =0.2). Raw amplitude data from dual cell recordings were analyzed using Wilcoxon signed rank test (p values indicated above). Normalized data were analyzed using a one-way ANOVA followed by the Mann−Whitney test. Scale bars: 30 ms, 50 pA.

### CaMKII binding to GluN2B is essential for its synaptic action

Biochemical studies indicate that CaMKII binds to GluN2B (Bayer et al., 2001; Bayer et al., 2006; O’Leary et al., 2011; Shen and Meyer, 1999; Strack and Colbran, 1998) and disrupting this binding impairs LTP (Barria and Malinow, 2005; Halt et al., 2012). To directly address the importance of this binding we overexpressed CA CaMKII under conditions that disrupt the binding of CA CaMKII to GluN2B (**Fig. 3A**). **Introducing** a mutant form of GluN2B (L1298A-R1300Q; GluN2B*), in which binding to the CaMKII surface groove is disabled (Halt *et al*., 2012), causes a depression in AMPAR responses after short OE (**Fig. 3B** and **D**, n =20). Short OE of both GluN2B* and CA CaMKII generates responses no larger than those recorded with GluN2B* alone (**Fig. 3C** and **D**, n =31), implying that CaMKII binding to GluN2B is required for its synaptic action.

**Figure 3.**
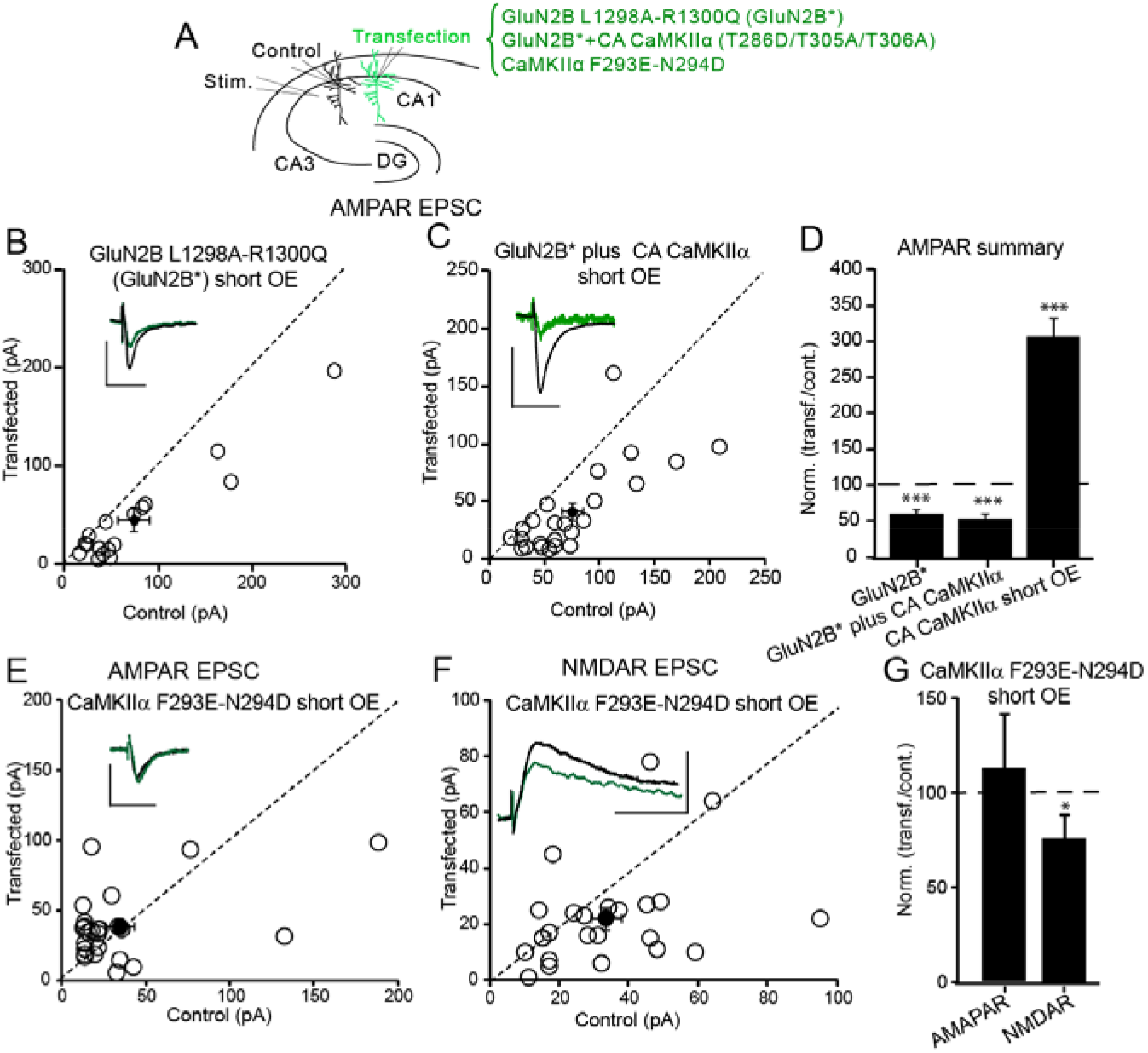
CaMKII binding to GluN2B is essential for its synaptic action. **A** Schematic diagram showing the electrophysiological approach and the transfected GluN2B and CaMKIIα mutate constructs (Control represents the wild type, un-transfected neurons). **B, C** Scatterplots showing amplitudes of AMPAR EPSCs for single pairs (open circles) of control cells and cells expressing GluN2B*(GluN2B L1298A-R1300Q) 4-5 days (short OE) (**B** = 20 pairs), and GluN2B*+ CA CaMKII 4-5 days (short OE) (**C**, n =31 pairs). Filled circles indicate mean ± SEM (**B**, Control = 73.4 ± 16.8; GluN2B*(GluN2B L1298A-R1300Q) short OE = 44.9 ± 11.6, p < 0.01; **C**, Control = 74.9 ± 9.4; GluN2B*+ CA CaMKII short OE = 41.5 ± 7.4, p < 0.01). **D** Bar graph of ratios normalized to control (%) summarizing the mean± SEM of AMPAR EPSCs (61.5 ± 7, p < 0.001). **E** Scatterplots showing amplitudes of AMPAR EPSCs for single pairs (open circles) of control cells and cells expressing CaMKII F293E-N294D for 1-5 days (short OE) (n = 25 pairs). Filled circle indicate mean ± SEM. (Control = 34.2 ± 8.9; CaMKII F293E-N294D short OE = 38.8 ± 8.8 p = 0.98). **F** Scatterplots showing amplitudes of NMDAR EPSCs for single pairs (open circles) of control and transfected cells of CaMKII F293E-N294D 1-5 days (short OE) (n = 25 pairs). Filled circles indicate mean ± SEM. (Control= 33.3 ± 4.4; CaMKII F293E-N294D short OE = 22.1 ± 3.9, p = 0.9). **G** Bar graph of ratios normalized to control (%) summarizing the mean ± SEM of APMAR and NMDAR EPSCs of values represented in **E** (113 ± 22, p =0.8) and **F** (77 ± 12.6, p <0.05). Raw amplitude data from dual cell recordings were analyzed using Wilcoxon signed rank test (p values indicated above). Normalized data were analyzed using a one-way ANOVA followed by the Mann−Whitney test. Scale bars: 30 ms, 50 pA

To further address the role of GluN2B binding in the action of CaMKII, we took advantage of a mutation in CaMKII (CaMKII (F293E/N294D)) that appears to have a conformation that is sufficiently open for substrates but not for GluN2B binding. This mutant even has a higher constitutive activity than CaMKII (T286D) (Yang and Schulman, 1999). However, just like wt CaMKII this mutation binds to GluN2B in the presence, but not in the absence of Ca^2+^/CaM (**Fig. S2G**). As expected, this mutant failed to enhance AMPAR responses (**Fig. 3E** and G, n = 25). And NMDAR responses were unchanged (**Fig. 3F** and **G**, n = 25). These findings indicate that, in the absence of GluN2B binding, constitutively active mutants fail to enhance synaptic responses.

The results discussed thus far examined the synaptic responses to the short-term expression of CaMKII (3-5 days). Long OE of CA CaMKII caused a profound depression in both the AMPAR (**Fig. S2A** and **S2C**, n =14) and NMDAR responses (**Fig. S2B** and **S2C**, n = 14). In addition, long OE of constitutively active CaMKII (F293E-N294D), which cannot bind to GluN2B (**Fig. S2G**), also causes a profound depression of both AMPAR (**Fig. S2D** and **S2F**, n =24) and NMDA responses (**Fig. S2E**, n = 24). Does this depression require kinase activity? While prolonged expression of “CA” CaMKII (D135N) enhances AMPAR responses as presented above (**Fig. S1A-C**), it had no effect on NMDA responses (**Fig. S1E**, n = 27). In addition, prolonged expression of CaMKII (F293E-N294D) with the kinase dead mutation (D135N) failed to depress either the AMPAR (**Fig. S2H**, n =24) or NMDAR response (**Fig. S3I**, n = 25). These results indicated that the deleterious effects on synaptic transmission after the prolonged expression of CA CaMKII is due to unrestricted phosphorylation of synaptic proteins.

### CaMKII autophosphorylation is required for LTP induction

Our results indicate that a kinase dead mutant of CaMKII held in an open configuration (T286D) is fully functional in potentiating synapses. The remaining question is whether any enzymatic activity is required for the action of CaMKII. In theory, the action of CaMKII in LTP could be purely structural: the interaction of Ca/CaM with CaMKII, by promoting the open state conformation, would enable the binding of the kinase to GluN2B, without the need of any enzymatic activity. Indeed, a recent study (Chang et al., 2017) reported normal LTP in the phosphonull mutant (T286A) knockin mouse. We therefore reexamined the role of phosphorylation in the induction of LTP using the pairing protocol of Chang et. al. (Chang *et al*., 2017) (pairing 40 Hz, 15 s with 0 mV depolarization). With this protocol, we observe an LTP that is similar in magnitude to that observed with our standard induction protocol (pairing 2 Hz, 90 s with 0 mV depolarization) (**Fig. 4A**, black circles). This LTP is largely blocked by the NMDAR blocker APV (**Fig. 4A**, red circles, n = 8) and by the double knockout of both CaMKIIα and CaMKIIβ (**Fig. 4A**, grey circles, n =10). We next replaced wt CaMKII with a kinase dead mutant (D135N). However, no LTP was recorded (**Fig. 4B**, yellow circles, n =11), in agreement with a previous study using the K42R kinase dead mutant knockin mouse (Yamagata et al., 2009). Finally, we examined LTP in cells expressing CaMKII (T286A) on a null background. We were unable to detect NMDAR-dependent LTP (**Fig. 5B**, green circles, n =10). The only difference between these experiments and those of Chang et al., is that they used T286A knockin mice, while we used *in utero* electroporation. It is unclear whether this difference explains the different results. Thus, our results indicated that CaMKII T286 phosphorylation is required for LTP induction.

**Figure 4.**
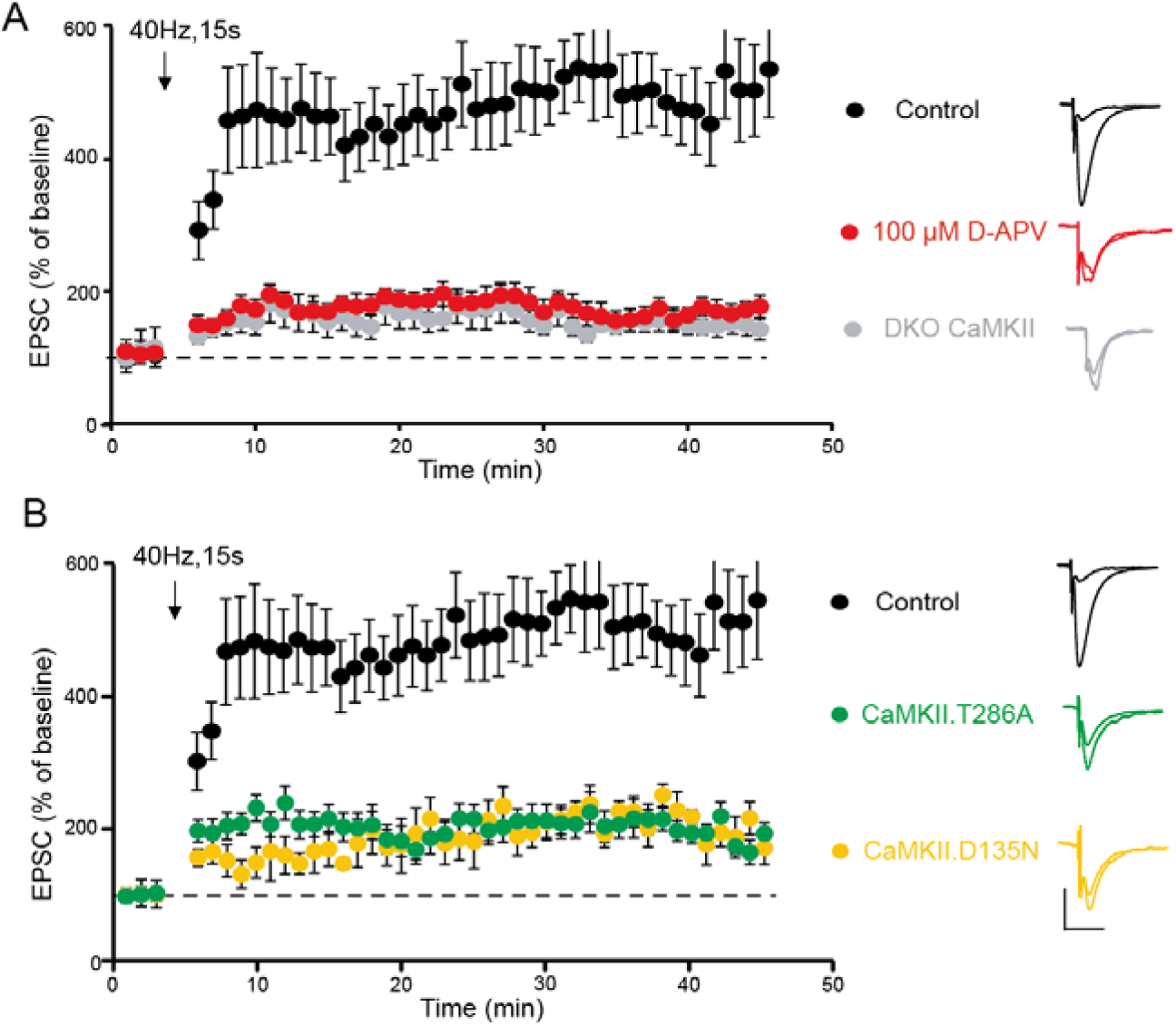
CaMKIIα T286A and CaMKII D135N fail to support LTP. **A** 40 Hz, 15s pairing protocol generates robust LTP in WT CA1 hippocampus cells (n = 13 control cells, black circles). LTP is largely blocked by 100 μM APV (Red circles, n = 8) and DKO CaMKII (CaMKIIα and CaMKIIβ) (Grey circles, n = 10, 6 simultaneous recordings included**). B** Replacement of CaMKIIα and CaMKIIβ with CaMKIIα D135N (Yellow circles, n =11, 7 simultaneous recordings included) or CaMKIIα T286A (green circles, n =10, 6 simultaneous recordings included) fail to support LTP. For all LTP graphs, control cells are shown as black filled circles ±SEM and transfected cells are shown as color circles ±SEM as defined. Traces show representative currents before and after LTP induction (scale bar: 50 pA/30 ms

**Figure 5.**
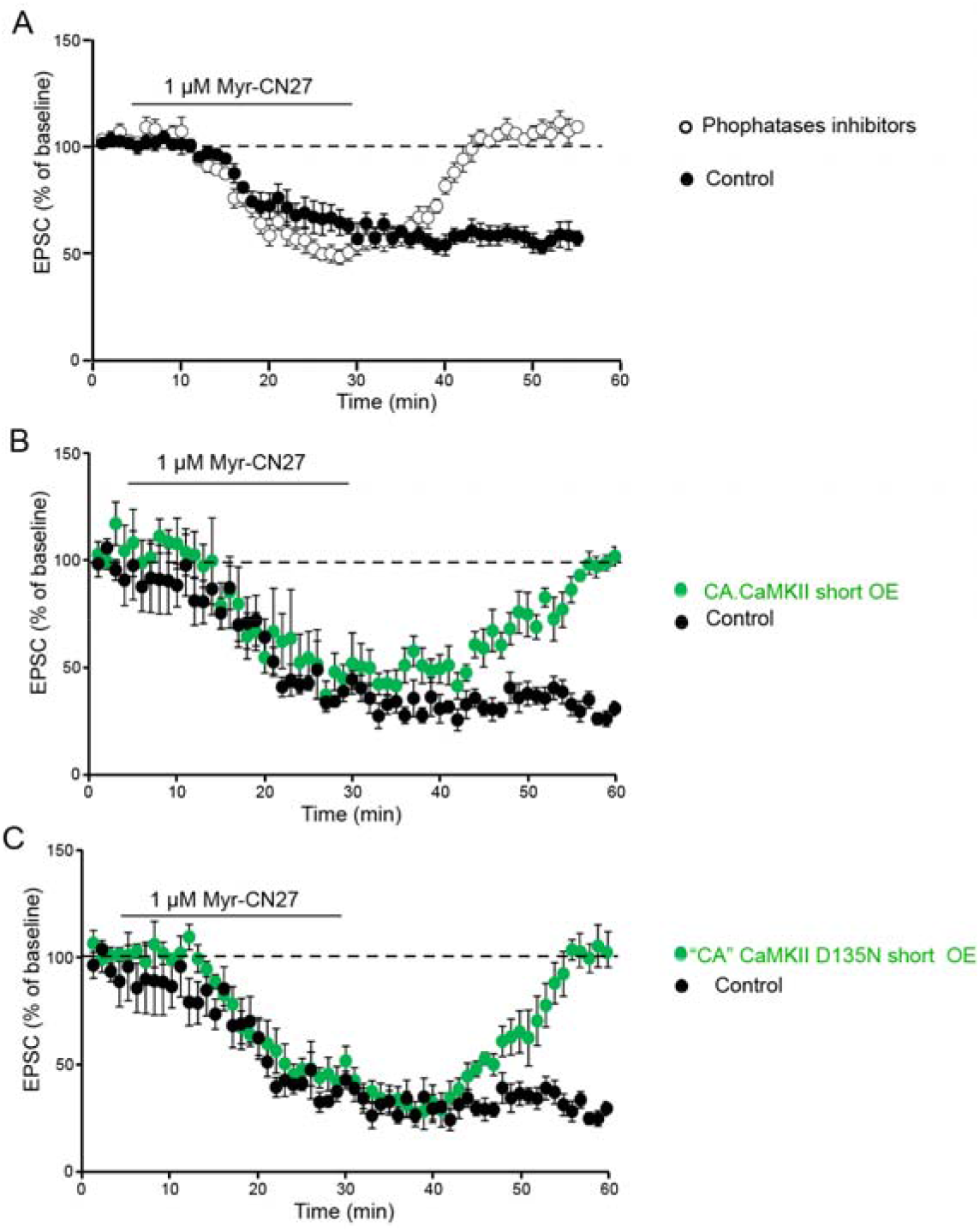
CaMKII T286 phosphorylation is required maintain CaMKII synaptic memory. **A** Plots show mean ± SEM of AMPAR EPSCs for transient application of 1μM Myr-CN27 (filled circles, n = 20). The experiment is repeated in the presence of phosphatases inhibitors (open circles, n = 23). **B** Plots show mean ± SEM AMPAR EPSC amplitude of control cells (black circles, n = 8) and cells transfected with CA CaMKII (green circles, n =8, 5 simultaneous recordings included). **C** Plots show mean ± SEM AMPAR EPSC amplitude of control cells (black circles, n = 8) and cells transfected with “CA” CaMKII D135N (green circles, n = 7, 6 simultaneous recordings included).

### CaMKII T286 autophosphorylation is required for maintaining synaptic memory

The results presented thus far indicate that T286 must be autophosphorylated for the induction of LTP. Is the autophosphorylation also required for the maintenance of LTP? To address this issue, we turned to the endogenous constitutive CaMKII component of synaptic transmission (Incontro *et al*., 2018; Lee *et al*., 2022; Sanhueza *et al*., 2011; Tao *et al*., 2021). We (Tao *et al*., 2021) and others (Sanhueza *et al*., 2011; Sanhueza and Lisman, 2013) have presented evidence that this constitutive CaMKII action, either enzymatic or binding to GluN2B, represents a CaMKII synaptic memory trace acquired prior to slice preparation. Does this trace require continued T286 autophosphorylation? The bath application of the CaMKII inhibitor myr-CN27 reverses this memory trace and importantly there is no recovery of synaptic transmission following the wash out of the inhibitor (**Fig. 5A**, black circles, n = 20) (Lee *et al*., 2022; Tao *et al*., 2021). It is important to keep in mind that CN27 has two actions: it blocks kinase activity (Chang et al., 1998; Pellicena and Schulman, 2014; Sanhueza et al., 2007; Vest et al., 2007) and it disrupts the binding of CaMKII to GluN2B (Ozden *et al*., 2022; Sanhueza *et al*., 2011; Vest *et al*., 2007). Thus, the depression caused during the myr-CN27 application is likely due to disruption of the binding of CaMKII to GluN2B. The failure of the responses to recover after washout of myr-CN27 would indicate that the release CaMKII is incapable of rebinding to GluN2B, perhaps due to its dephosphorylation. If this is the case one would predict that repeating this experiment in the presence of phosphatase inhibitors, should allow for the rebinding of the autophosphorylated CaMKII to rebind to GluN2B.

Phosphatase inhibitors were either bath applied (calyculin A, n =11) or added to the pipette solution (microcystin LR, n =12) Since both manipulations gave the same result, we have combined the data. The inhibitors have no effect on the magnitude of the synaptic depression during the myr-CN27 application (**Fig. 5A**, open circles). However, following the washout of myr-CN27 the responses rapidly recover in the presence of phosphatase inhibitors (**Fig. 5A**, open circles). These findings suggest that the acute inhibition caused by myr-CN27 is due to the dissociation of CaMKII from GluN2B and not the dephosphorylation of T286. To test this idea, we repeated the experiment in neurons expressing either CA CaMKII (**Fig. 5B**, green circles, n = 7) or the “CA” CaMKII (D135N) (**Fig. 5C** green circles, n = 7). The responses in neurons expressing either CA CaMKII or “CA” CaMKII (D135N) rapidly recovered. In contrast, as expected, in either simultaneously recorded control cells (n = 15) or interleaved cells (n = 11) there was no recovery following the transient application of myr-CN27. Taken together these results indicate that the depression in synaptic transmission during the application of myr-CN27 is due to the dissociation of active CaMKII from GluN2B, while the persistent depression following wash out of the inhibitor is due to the dephosphorylation of T286 and its failure to rebind to GluN2B. CA CaMKII is phosphatase resistant and GluN2B binding enabled because of its open state but can be displaced by myr-CN27. “CA” CaMKII (D135N) is also phosphatase resistant, catalytic activity independent and GluN2B binding enabled, but can also be displaced by myr-CN27. This further confirms that no phosphorylation is necessary for recovery. These results show that the maintenance of LTP requires T286 autophosphorylation and that binding of CaMKII to GluN2B shields this phosphorylation from phosphatases.

## Discussion

Biochemical studies have established that CaMKII has two enzymatic actions, autophosphorylation of T286, which maintains the enzyme in an open active state and phosphorylation of downstream targets, which is proposed to enhance synaptic transmission. However, the high levels of CaMKII in the PSD, ranging up to 10-30% (Chen et al., 2005; Kennedy et al., 1983; Peng et al., 2004) have long raised the possibility of a structural role for CaMKII (Kim et al., 2016; Lisman et al., 2002). We have used 6 different mutant forms of CaMKII to distinguish between the enzymatic and structural roles in its synaptic enhancing action and in LTP. There are two major conclusions (**Fig. S3**). First, we conclude that the autophosphorylation of T286 and the binding of CaMKII to GluN2B are essential for the synaptic enhancing action of CaMKII. The fact that no potentiation was seen immediately after LTP induction with either the kinase dead CaMKII or CaMKII (T286A), which are expected to bind to GluN2B (Bayer *et al*., 2001; Bayer *et al*., 2006; O’Leary *et al*., 2011; Shen and Meyer, 1999; Strack and Colbran, 1998), indicates that this binding is rapidly reversible or that it does not occur under these conditions. The essential role of CaMKII binding to GluN2B emphasizes the central role of the NMDAR in LTP; it is initiates LTP by increasing spine Ca^2+^ levels and it maintains LTP by its binding GluN2B. T286 autophosphorylation of CaMKII is required for stabilizing the CaMKII/GluN2B complex. This finding nicely compliments the recent phase separation studies of Hosokawa et al. (Hosokawa et al., 2021) showing that while the condensates formed by Ca^2+^/CaM-bound CaMKII and GluN2B do not require T286 phosphorylation, it is required for the condensates to remain intact following the removal of Ca^2+^/CaM.

The second major conclusion is that phosphorylation of the many identified downstream synaptic target proteins of CaMKII, such as GluA1, TARPs, synGAP, RhoGEFs, PSD-95 etc., that have long been implicated in LTP (reviewed in (Nicoll and Schulman, 2023)), is dispensable for its synaptic enhancement, thus establishing a structural role of CaMKII. In fact, the prolonged phosphorylation of one or more synaptic proteins by CA CaMKII had a profound deleterious effect on both AMPAR and NMDAR responses. This finding implies that during the maintenance of LTP the active CaMKII is prevented access to these target proteins by its sequestered binding to GluN2B. What role these target proteins play in the action of CaMKII requires further investigation. The present results focus attention on the CaMKII/GluN2B complex as a structural hub. Again, the recent studies of Hosokawa et al. (Hosokawa *et al*., 2021) and Cai et al. (Cai et al., 2021) are informative. The authors show that CaMKII and GluN2B undergo phase separation forming lasting condensates after the removal of Ca^2+^/CaM. However, phosphatase can rapidly remove phosphate from T286 and causing the rapid disassembly of CaMKII/GluN2B condensates. Our results showing that dephosphorylation of T286 prevents the enhancing action of CaMKII, provides physiological validation for these phase separation results. Furthermore, the addition of PSD-95 and TARPs, a proxy for AMPARs, results in a distinct phase in phase separation. Except for T286 phosphorylation, this molecular assembly is independent of phosphorylation. It will be of interest to establish whether the phase separation so clearly established with the synaptic component proteins in solution can also explain our present results with intact synapses.

Finally, our study reveals that the enzymatic activity of synaptic CaMKII is dedicated to the regulation of its open state via autophosphorylation, thereby allowing this extremely abundant enzyme to function as a synaptic scaffold protein. Since autophosphorylation and consequent conformational change is a rather general property of protein kinases, it will be interesting to explore whether other protein kinases also adopt similar autophosphorylation-regulated structural roles for cellular processes.

## Acknowledgements

We thank Dr. M. Scanziani for his extensive comments on the manuscript. This work was supported by a grant from National Science Foundation of China (82188101), the Minister of Science and Technology of China (2019YFA0508402), an HFSP Research Grant (RGP0020/2019), and grants from Research Grant Council of Hong Kong (AoE-M09-12, 16104518, and 16101419) to M.Z. and a grant from NIH (R01MH117139) to R.A.N.

## Methods

### DNA constructs and chemical agents

The cDNA sequences of rat CaMKIIα (Uniprot: P11275) were PCR-amplified from rat brain cDNA library. CaMKII DKO and GluN2B mutated constructs have been described previous (Incontro et al., 2018). Myr-CN27 (Calbiochem. Inc. catalog # 208921), microcystin LR (Sigma-Aldrich. catalog # 475823) and calyculin A (VWR. catalog # EI192) were purchased. For the constitutively active CaMKII we used the triple mutant CaMKII (T286D/T305A/T306A) (Incontro *et al*., 2018; Pi *et al*., 2010), referred to as CA CaMKII. T305 and T306 are phosphorylation sites within the CaM-binding domain that become autophosphorylated by an autonomous kinase when Ca^2+^/CaM dissociates, blocking any rebinding; the phosphonull mutations T305A/T306A are used to avoid that complication.

### Protein Expression and Purification

Wild type and D135N mutant of CaMKIIα holoenzymes were expressed and purified according to the previous reported protocols (Cai et al., 2023; Cai *et al*., 2021; Chao et al., 2010). Briefly, variants of CaMKII with N-terminal His_6_-SUMO tag were co-expressed with λ phosphatase in *Escherichia coli* BL21-CodonPlus(DE3)-RIL cells (Agilent Technologies) in LB medium at 16 □ for 24 h. The recombinant CaMKII proteins were extracted by high pressure homogenizer and purified by Ni^2+^-NTA Sepharose 6 Fast Flow resin (Cytiva) and Superose 6 26/600 (Cytiva) gel filtration chromatography. After the His_6_-SUMO tag was removed by Ulp1 protease, proteins were further purified by MonoQ anion exchange chromatography and Superose 6 10/300 gel filtration chromatography. The final column buffer and well as the storage buffer was 50 mM Tris pH 8.0, 200 mM NaCl, 10% glycerol, 5 mM DTT. All other proteins were expressed in *Escherichia coli* BL21-CodonPlus(DE3)-RIL cells (Agilent Technologies) in LB medium at 16 □ overnight. Recombinant proteins were firstly purified using Ni^2+^-NTA Sepharose 6 Fast Flow affinity chromatography (Cytiva) for His_6_-tagged proteins or Glutathione Sepharose 4 Fast Flow affinity chromatography (Cytiva) for GST-tagged proteins and followed by a step of gel filtration chromatography using Superdex 200 26/600 or Superdex 75 26/600 column (Cytiva).

### GST Pull Down Assay

GST pull down assay was carried out as previously described (Cai *et al*., 2023; Cai *et al*., 2021). Briefly, HEK293T cells were transiently transfected with various GFP-CaMKII encoding plasmids using PEI, then harvested at 20 h post transfection and lysed using the lysis buffer containing 50 mM Tris pH 7.5, 150 mM NaCl, 1% Triton X-100 and a cocktail of protease inhibitors (Merck). The mixture of lysate and GST-fused GluN2B (a.a. 1259-1310) was incubated at 4 □ for 30 min. After centrifugation at 16,873 g for 5 min, the supernatant was mixed with 40 μL of fresh Glutathione Sepharose resin and incubated for another 30 min. After extensive washing, the captured proteins were eluted by SDS-PAGE loading buffer by boiling, resolved by SDS-PAGE, and immunoblotted with specific antibodies. Protein signals were visualized by an HRP-conjugated secondary antibody (Jackson ImmunoResearch) and western HRP substrate (Sangon).

### CaMKII kinase activity assay using phos-tag SDS-PAGE

15% acrylamide gel with 25 μM phos-tag acrylamide (AAL-107, Wako) and 250 μM MnCl_2_ were prepared for phos-tag SDS-PAGE. N-terminal Trx-tagged Syntide-2 was purified and used as the substrate for CaMKII kinase activity assay. The experiments were carried out in the buffer containing 50 mM Tris pH 7.5, 150 mM NaCl, 1 mM CaCl_2_, 10 mM MgCl_2_, 0.5 mM ATP, 3 μM CaM, 1 μM CaMKII and 100 μM Trx-Syntide-2. The reactions were quenched by 2×SDS loading buffer at different time points.

### Lentivirus Production

Three T-75 flasks of rapidly dividing HEK293T cells (ATCC) were transfected with 27 mg FUGW-CaMKII 2gRNA-CaMKII variant-mCherry, plus helper plasmids pVSV-G (18 mg) and psPAX2 (27 mg) using FuGENE HD (Promega). DNA was incubated with 210 ml FuGENE HD in 4.5 ml Opti-MEM (Life Technologies) before transfection, according to the manufacturer’s directions. Forty hours and then again at 3 days supernatant was collected, filtered, and concentrated using the PEG-it Virus Precipitation Solution (System Biosciences) according to the manufacturer’s directions. The resulting pellet was re-suspended in 150 ml Opti-MEM, flash-frozen with dry ice, and stored at -80°C.

### P0 injection

Rosa26-Cas9 knock-in mice were used. The mice of both sexes were purchased from Jackson labs (Stock No. 026179). These mice constitutively express Cas9 and GFP. Concentrated lentiviruses or AAV virus (Made by Virovek) were injected bilaterally into the medial hippocampi of hypothermia anesthetized P0-P1 pups, by free hand. 2 μl per hemisphere was slowly delivered. All experiments were performed in accordance with established protocols approved by the University of California San Francisco Institutional Animal Care and Use Committee.

### In utero electroporation

In utero electroporation was performed as previously described (Quirogo et al., 2007). E14 pregnant WT (CD-1) mice were anesthetized with 2% isoflurane. Embryos were exposed and injected with∼1.5 μl of mixed plasmid DNA with Fast Green (Sigma Aldrich) into the lateral ventricle via a beveled micropipette. Each embryo was electroporated with five 27 V pulses of 50 ms, delivered at 1 Hz, using platinum tweezer-trodes in a square-wave pulse generator (BTX Harvard Apparatus). The positive electrode was placed in the lower right hemisphere and the negative electrode placed in the upper left hemisphere. Following electroporation, the embryos were placed back into the abdominal cavity and abdominal muscle and skin were sutured. All experiments were performed in accordance with established protocols approved by the University of California, San Francisco’s Institutional Animal Care and Use Committee.

### Acute slice preparation

Acute hippocampal slices were prepared from P18-P28 mice. Mice were anesthetized with isoflurane. Brains were removed and sliced into 300 μm near-horizontal sections using Microslicer DTK-Zero1 (Ted Pella). Slices were then transferred to a holding chamber containing ACSF (in mM) (125 NaCl, 2.5 KCl, 1.25 NaH_2_PO_4_, 25 NaHCO_3_, 11 glucose, 1 MgSO_4_, 2 CaCl_2_ saturated with 95% O_2_/5% CO_2_) and incubated for 20 minutes at 37°C and then kept at room temperature until use.

### Slice culture and transfection

Hippocampal organotypic slice cultures were prepared from 7-9 day old rats as previously described (Stoppini et al., 1991). Transfections were carried out 48 h after. Briefly, 50 μg DNA was coated on 1 μm diameter gold particles in 0.5 mM spermidine, precipitated with 0.1mM CaCl_2_, and washed four times in pure ethanol. The gold particles were coated onto PVC tubing, dried using ultra-pure N_2_ gas, and stored at 4 °C in desiccant. DNA-coated gold particles were delivered with a Helios Gene Gun (BioRad). Slices were maintained at 34°C with media changes every two days.

### Electrophysiological recording

All electrophysiological recordings were carried out on an upright Olympus BX51WI microscope and collected using a Multiclamp 700B amplifier (Molecular Devices). During recording, slices were maintained in ACSF (in mM): 125 NaCl, 2.5 KCl, 1.25 NaH_2_PO_4_, 25 NaHCO_3_, 11 glucose saturated with 95% O_2_; 5% CO_2_) containing 1 MgSO_4_, 2 CaCl_2_ during acute slice recordings and 4 MgSO_4_, 4 CaCl_2_ during slice culture recordings. Transfected cells were identified visually using fluorescence and recorded simultaneously with a neighboring control cell. All recordings were carried out at 20-25 °C using glass patch electrodes filled with an intracellular solution (in mM): 135 CsMeSO_3_, 10 HEPES, 8 NaCl, 0.3 EGTA, 4 Mg-ATP, 0.3 Na-GTP, 5 QX-314, and 0.1 spermine). Synaptic currents were elicited by stimulation of the Schaffer collaterals with a bipolar electrode (Micro Probes). AMPAR- and NMDAR-mediated responses were collected in the presence of 100 μM picrotoxin to block inhibition. 5 μM 2- Chloroadenosine was used to suppress epileptic activity in slice culture. The bipolar stimulating electrode was placed in stratum radiatum. AMPAR EPSCs were evoked while voltage clamping cells at -70 mV, and the amplitude was determined by measuring the peak of this response. NMDAR EPSCs were obtained while voltage clamping cells at +40 mV and measured at 100 ms. Series resistances typically ranged from 10 to 20 MΩ; a cell pair was discarded if the series resistance of either cell increased to >30 MΩ. Statistical difference was determined using a two-tailed paired t test.

LTP was induced via a pairing protocol of 40 Hz stimulation for 15 s at a holding potential of 0 mV, after recording a 3-5 min baseline, but not more than 6 min after breaking into the cell. All LTP experiments were carried out in acute slices. Simultaneous dual whole-cell recordings were made in a transfected CA1 pyramidal cell and a neighboring wild-type cell. In some cases, one of the paired cells was lost during the experiment, then the recordings were considered until that point. In cases where one cell was lost the remaining cell was considered for the averages.

### Quantification and statistical analysis

Statistical parameters including the definitions and exact values of n (e.g. number of experiments, number of droplets), distributions and deviations are reported in the Figures and corresponding Figure Legends. Statistical analysis was performed in GraphPad Prism.

