## supplemental data for "CaMKII autophosphorylation but not downstream kinase activity is required for synaptic memory"

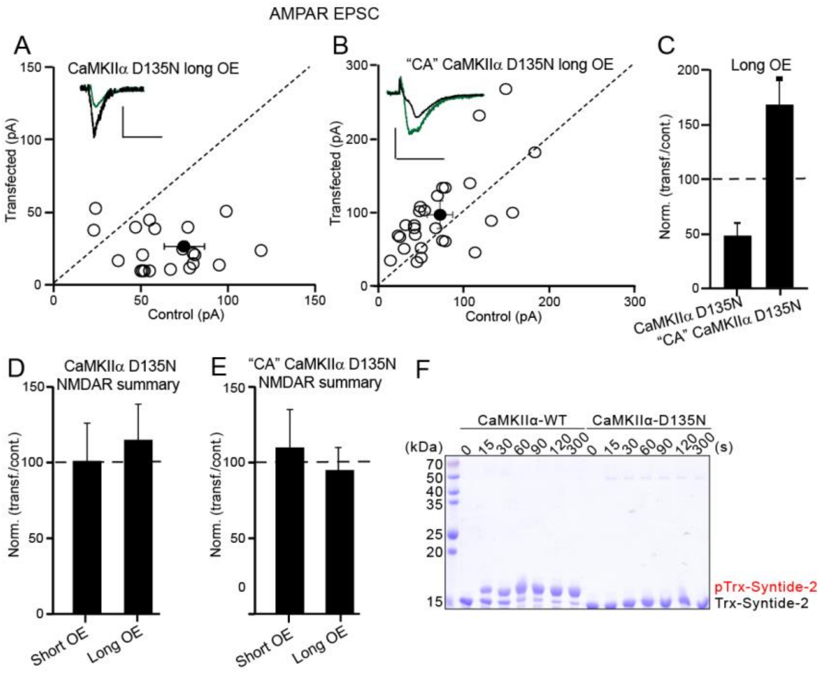


**Figure S1. Kinase dead mutations CaMKIIα enhances synaptic transmission for overexpression in present with GluN2B binding.**

**A, B** Scatterplots showing amplitudes of AMPAR EPSCs for single pairs (open circles) of control and overexpressing cells of CaMKII D135N 14 days (long OE) (CaMKII D135N replacement) (**A**, n = 25 pairs), and "CA" CaMKIIα D135N 14 days (long OE) (CA CaMKII D135N replacement) (**B**, n = 20 pairs). Filled circle indicate mean ± SEM. (**A**, Control = 74.7 ± 11.7; CaMKII D135N long OE = 26.8 ± 3.6 $p < 0.001$; **B**, Control = 72.1 ± 14.8; "CA" CaMKII D135N 14 days (long OE) = 98.2 ± 19 $p < 0.01$). **C** Bar graph of ratios normalized to control (%) summarizing the mean ± SEM of AMPAR EPSCs of **A** (49 ± 10, $p < 0.001$) and **B** (170 ± 21, $p < 0.005$). **D** Bar graph of ratios normalized to control (%) summarizing the mean ± SEM of NMDAR EPSCs of CaMKII D135N 2-4 days (short OE) (101 ± 25, p =0.6) and 14 days (long OE) (110 ± 24, p =0.8). **E** Bar graph of ratios normalized to control (%) summarizing the mean ± SEM of NMDAR EPSCs of "CA" CaMKII D135N 2-4 days (short OE) (115 ± 30, p =0.9) and 14 days (long OE) (95 ± 15, p =0.7). Raw amplitude data from dual cell recordings were analyzed using Wilcoxon signed rank test (p values indicated above). Normalized data were analyzed using a one-way ANOVA followed by the Mann−Whitney test. Scale bars: 30 ms, 50 pA.

Fig S2

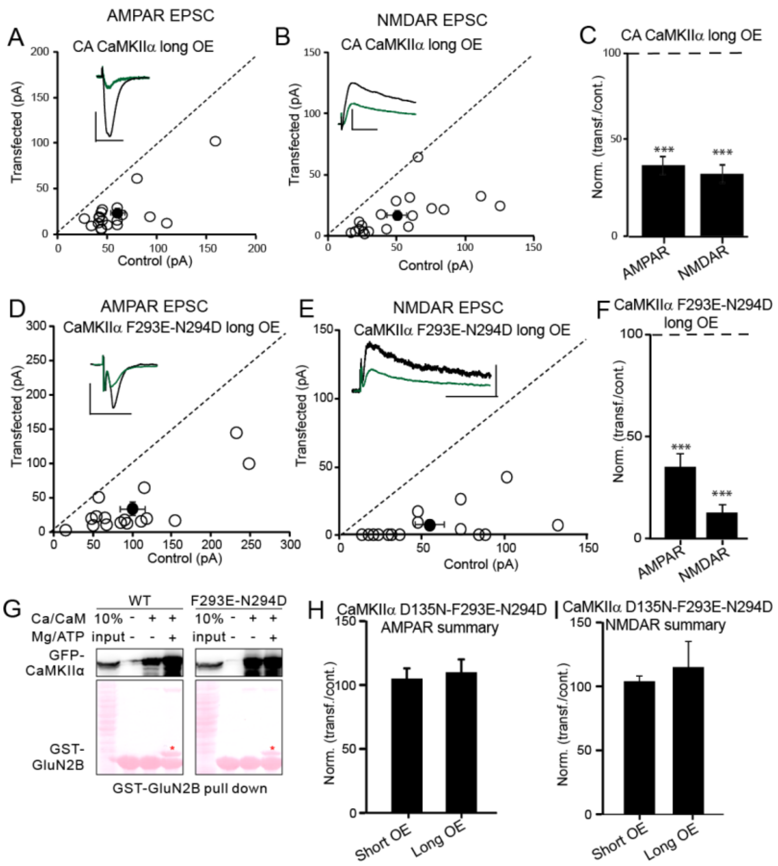


**Fig. S2 Prolonged expression of CA CaMKII depress synaptic transmission**

**A** Scatterplots showing amplitudes of AMPAR EPSCs for single pairs (open circles) of control and overexpressing cells of CA CaMKIIα (CaMKIIα T286D-T305A-T306A ) 10 days (long OE) (n = 14 pairs). Filled circle indicate mean ± SEM. (Control = 61.3 ± 7.2; CA CaMKIIα long OE = 23.7 ± 5.2 p < 0.0001). **B** Scatterplots showing amplitudes of NMDAR EPSCs for single pairs (open circles) of control and transfected cells of CA CaMKIIα 10 days (long OE) ( n = 14 pairs). Filled circles indicate mean ± SEM. (Control = 50 ± 7.2; CA CaMKIIα 10 days = 16.9 ± 3.6, p < 0.001). **C** Bar graph of ratios normalized to control (%) summarizing the mean ± SEM of AMPAR and NMDAR EPSCs of values represented in **A** (38.5 ± 4.8, p < 0.0001) and **B** (32.6 ± 5, p < 0.0001).

**D** Scatterplots showing amplitudes of AMPAR EPSCs for single pairs (open circles) of control and overexpressing cells of CaMKIIα F293E-N294D 10 days (long OE) ( n = 24 pairs). Filled circle indicate mean ± SEM. (Control = 100.1 ± 16; CaMKIIα F293E-N294D long OE = 34.2 ± 9.7 p < 0.001). **E** Scatterplots showing amplitudes of NMDAR EPSCs for single pairs (open circles) of control and transfected cells of CaMKIIα F293E-N294D10 days (n = 24 pairs). Filled circles indicate mean ± SEM. (Control= 54.3 ± 8.9; CaMKIIα F293E-N294D 10 days )long OE) = 8 ± 3.1). **F** Bar graph of ratios normalized to control (%) summarizing the mean ± SEM of AMPAR and NMDAR EPSCs of values represented in D (32 ± 5.5, $p < 0.0001$) and **E** (12.4 ± 3.7, $p < 0.0001$). **G** GST-GluN2B pull-down. In the presence, but not in the absence of $Ca^{2+}$/CaM, CaMKIIα wt and CaMKIIα F293E-N294D bind to GluN2B. **H** Bar graph of ratios normalized to control (%) summarizing the mean ± SEM of AMPAR EPSCs of CaMKIIα D135N-F293E-N294D 1-5 days (short OE) (105 ± 8, p =0.7) and 10 days (long OE) (110 ± 18, p =0.8). **I** Bar graph of ratios normalized to control (%) summarizing the mean ± SEM of NMDAR EPSCs of CaMKIIα D135N-F293E-N294D 1-5 days (short OE) (104 ± 5, p =0.9) and 10 days (long OE) (115 ± 20, p =0.7). Raw amplitude data from dual cell recordings were analyzed using Wilcoxon signed rank test (p values indicated above). Normalized data were analyzed using a one-way ANOVA followed by the Mann−Whitney test. Scale bars: 30 ms, 50 pA

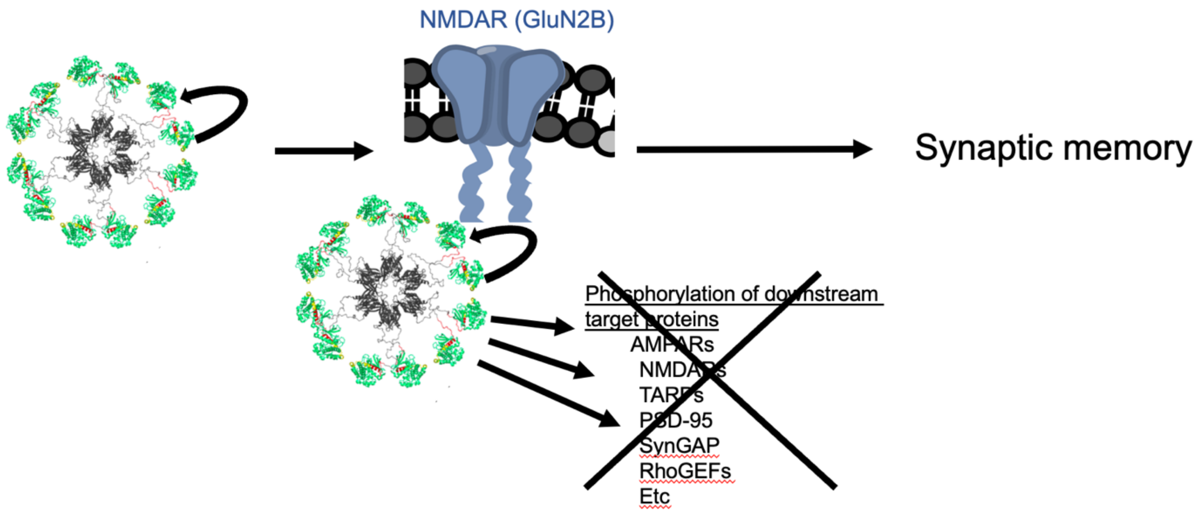


**Fig. S3 Autophosphorylation of CaMKII T286 is required for synaptic memory, but not phosphorylation of downstream synaptic proteins.** Autophosphorylation of CaMKII T286 is required for synaptic memory by stabilizing the binding of CaMKII to GluN2B. The numerous downstream synaptic protein targets of CaMKII are not required for synaptic memory. Rather the CaMKII/GluN2B complex serves as a structural signaling hub to enhance and maintain synaptic strength.
